# Thousands of previously unknown phages discovered in whole-community human gut metagenomes

**DOI:** 10.1101/2020.10.07.330464

**Authors:** Sean Benler, Natalya Yutin, Dmitry Antipov, Mikhail Raykov, Sergey Shmakov, Ayal B. Gussow, Pavel Pevzner, Eugene V. Koonin

## Abstract

**Background:** Double-stranded DNA bacteriophages (dsDNA phages) play pivotal roles in structuring human gut microbiomes; yet, the gut phageome is far from being fully characterized, and additional groups of phages, including highly abundant ones, continue to be discovered by metagenome mining. A multilevel framework for taxonomic classification of viruses was recently adopted, facilitating the classification of phages into evolutionary informative taxonomic units based on hallmark genes. Together with advanced approaches for sequence assembly and powerful methods of sequence analysis, this revised framework offers the opportunity to discover and classify unknown phage taxa in the human gut.

**Results:** A search of human gut metagenomes for circular contigs encoding phage hallmark genes resulted in the identification of 3,738 apparently complete phage genomes that represent 451 putative genera. Several of these phage genera are only distantly related to previously identified phages and are likely to found new families. Two of the candidate families, “Flandersviridae” and “Quimbyviridae”, include some of the most common and abundant members of the human gut virome that infect *Bacteroides, Parabacteroides* and *Prevotella*. The third proposed family, “Gratiaviridae”, consists of less abundant phages that are distantly related to the families *Autographiviridae*, *Drexlerviridae* and *Chaseviridae*. Analysis of CRISPR spacers indicates that phages of all three putative families infect bacteria of the phylum Bacteroidetes. Comparative genomic analysis of the three candidate phage families revealed features without precedent in phage genomes. Some “Quimbyviridae” phages possess Diversity-Generating Retroelements (DGRs) that generate hypervariable target genes nested within defense-related genes, whereas the previously known targets of phage-encoded DGRs are structural genes. Several “Flandersviridae” phages encode enzymes of the isoprenoid pathway, a lipid biosynthesis pathway that so far has not been known to be manipulated by phages. The “Gratiaviridae” phages encode a HipA-family protein kinase and glycosyltransferase, suggesting these phages modify the host cell wall, preventing superinfection by other phages. Hundreds of phages in these three and other families are shown to encode catalases and iron-sequestering enzymes that can be predicted to enhance cellular tolerance to reactive oxygen species.

**Conclusions:** Analysis of phage genomes identified in whole-community human gut metagenomes resulted in the delineation of at least three new candidate families of *Caudovirales* and revealed diverse putative mechanisms underlying phage-host interactions in the human gut. Addition of these phylogenetically classified, diverse and distinct phages to public databases will facilitate taxonomic decomposition and functional characterization of human gut viromes.

## Background

The bulk of the human-associated virome resides in the distal gastrointestinal tract and is composed of tailed double-stranded (ds) DNA bacteriophages (dsDNA phages) [1–3] that, in the recent virus megataxonomy, are classified as the class *Caudoviricetes* under the phylum *Uroviricota* [4]. The ternary interactions between phages, bacteria and their human hosts are being elucidated at an increasing pace through experiments on model systems and sequencing of the uncultured community of viruses (virome) [5–9]. Comparisons of the human gut virome within and between individuals unveils remarkable longitudinal stability and high diversity of resident phages [2, 10, 11]. Although the human gut offers a rich source of phage genomic diversity, the virome so far has been explored to a much lesser extent than the whole community (metagenome), composed of viruses, Bacteria and Archaea. The rapid growth of the public whole-community metagenomic data offers the opportunity to identify numerous novel phage genomes lurking in metagenomes.

Depending on the terminal genomic arrangement, many complete phage genomes assemble into a ‘circular’ contig (i.e., a contig with direct terminal repeats) [12]. Thus, circularity can be used as one feature to identify putative complete phage genomes in viromes and metagenomes. However, the comparatively small size of dsDNA phage genomes (~50 kb, on average) [13] and the estimated low virus-to-microbe ratio in the gut (1: 10) [1] jointly translate into a relatively small amount of phage DNA present in whole community metagenomic libraries [14]. Moreover, similar-sized plasmids also assemble into circular contigs [15]. A recently developed computational method aims to address this problem by focusing specifically on the assembly of circular phage genomes and their automatic discrimination from plasmids based on gene content [16]. The genetic repertoire shared between plasmids and phages, for example, the *parABS* partitioning system encoded by both *Escherichia coli* phage P1 and plasmids [17], can obfuscate their automatic annotation-based discrimination and necessitate manual curation. Despite these challenges, there is a pressing need to reduce the amount of viral “dark matter” in the human gut by identifying and classifying phages for reference-based analyses [18, 19].

The global organization of the virosphere was recently captured in a comprehensive, unified framework using protein domains encoded by viral hallmark genes to infer evolutionary connections between major groups of viruses [4] and subsequently approved by the International Committee on the Taxonomy of Viruses (ICTV) as the comprehensive, multi-rank taxonomy of viruses. In particular, dsDNA viruses possess either the HK97 fold or the double jelly-roll fold in their major capsid proteins, along with distinct ATPases involved in capsid maturation, and thus appear to have independent origins, justifying their separation into two realms (the highest virus taxon rank) [4]. Tailed dsDNA phages, with their HK97 major capsid proteins, comprise the order *Caudovirales* within the class *Caudoviricetes*, under the phylum *Uroviricota* (that also include the distantly related herpesviruses of animals) and are further classified into 9 families. With the now formally recognized ability to classify viruses from sequence data alone [20], phylogenomic analysis of uncultured phage genomes can delineate novel taxa.

Here, we describe 3,738 completely assembled phage genomes discovered by analysis of 5,742 whole-community human gut metagenomes. Using abundance, taxonomy and genomic composition as criteria to select genomes for further scrutiny, three groups of phages, all infecting bacteria of the phylum Bacteroidetes comprising potential new families, were analyzed in detail. All these candidate families, named “Quimbyviridae”, “Flandersviridae”, and “Gratiaviridae” consist of phages infecting bacteria of the phylum Bacteroidetes, and the first two are widely distributed and abundant in human gut viromes. The phages in these families and others yet to be classified encode enzymes that are involved in the response of cells to oxidative stress, implicating phages in the tolerance of anaerobes to oxygen. Furthermore, comparative genomic analysis exposed genetic cassettes that are unique to some genera in each family and thus appear to be relatively recent acquisitions involved in phage-host interactions. Addition of all the phage genomes identified here to public databases will substantially expand the known phage diversity and augment taxonomic classification of the human gut virome.

## Methods

### Identification of phage genomes in human gut metagenomes

5,742 whole-community metagenome assemblies generated from human fecal samples were downloaded from the NCBI Assembly database (accessed 8/2019). To limit the search space to likely complete genomes, 95,663 ‘circular’ contigs (50-200 bp direct overlap at contig ends) were extracted from these assemblies. Next, 304 phage-specific protein alignments from the CDD database [21] and 117 custom alignments (Yutin et al., *in press*) were converted to Hidden Markov Models (HMMs) using hmmpress (v. 3.2.1). Proteins in the 95,663 contigs were predicted by Prodigal (v. 2.6.3) [22] in the metagenomic mode and searched against the set of 304 phage-specific HMMs using hmmsearch, with the relaxed e-value cutoff of < 0.05. Contigs with at least one hit (n = 4,907) were selected for a second round of searches after correcting for re-assigned codons, as follows. All contigs were searched for the presence of tRNAs using tRNA-scan-SE (v. 2.0) [23]. In 212 contigs, an amber stop codon-suppressor tRNA was identified. ORFs were re-predicted for these contigs with the amber stop codon re-assigned to glutamine, given that this reassignment is most commonly observed in human gut phages [24, 25]. The re-translated contigs were added back to the database and all contigs were subjected to a second profile search with a stricter e-value cutoff (< 0.01). Contigs were classified as phages when exceeding 3 kbp in length and possessing at least one ORF that matched a capsid, portal or large terminase subunit protein profile below the e-value threshold. The phage classifications were cross-checked with Seeker [26] and ViralVerify [16]. In cases where both tools classified a contig as non-phage, the protein annotations were examined manually, revealing four contigs of ambiguous identity that were discarded.

### Collection of phage genomes in GenBank

Taxonomic accession codes corresponding to all prokaryotic viruses were collected from the NCBI Taxonomy database and used to extract sequences longer than 3 kbp from the non-redundant nucleotide database (accessed 09/2019). The protein predictions for each genome sequence were retrieved using the ‘efetch’ functionality in the entrez direct command line tools [27]. Genomic sequences lacking protein predictions were discarded.

### Dereplication and annotation of phage genomes

The collections of GenBank and human gut phage genomes were each dereplicated at 95% average nucleotide identity across 80% of the genome length using dRep (v. 2.6.2) [28] and its associated dependencies, Mash [29] and FastANI [30], with all other settings left as default. The proteins from these contigs were collected and clustered at 95% amino acid identity across 33% of the protein length using mmclust [31]. The representative protein sequences were combined into a single BLAST database and compared against the multiple sequence alignments (MSAs) in the CDD database [21] with PSI-BLAST [32] at an evalue cutoff of 0.01. If the representative protein sequence produced a significant result, the representative and all constituent members of the protein cluster were annotated using the best hit.

### Phylogenetic reconstruction

Alignments of the large terminase subunit (TerL), capsid or portal protein were constructed as previously described [33]. Marker proteins from the metagenomic phages were combined with markers from GenBank phages into a single database and initially clustered to 50% amino acid identity using mmclust [31]. The clusters were aligned using MUSCLE [34]; cluster alignments were then compared to each other using HHsearch (v. 3.0) [35]. The cluster-cluster similarity scores were converted to distances as -ln(*S*_A,B_/min(*S*_A,A_,*S*_B,B_)), where S_A,B_ is similarity between the profiles A and B, then, an unweighted pair group method with arithmetic mean (UPGMA) dendrogram was constructed using the estimated cluster distances. Tips of the tree (depth <1.5) were used to guide the pairwise alignment of the clusters at the tree leaves with HHalign, creating larger protein clusters. The resulting alignments were filtered to remove sites with more than 50% gaps and a homogeneity lower than 0.1 [36]. The filtered alignment was used to construct an approximate maximum-likelihood tree using FastTree [37], with the Whelan-Goldman models of amino acid evolution and gamma-distributed site rates. Examination of the trees identified 353 nearly identical PhiX-174 sequences that were removed from subsequent analyses as a contamination from a sequencing reagent.

### Phage genome analysis

A gene-sharing network of phage genomes was constructed using Vcontact2 (v. 0.9.19) [38], with default search settings against the database of dereplicated GenBank phage genomes. The results were imported into Cytoscape (v. 3.8) [39] for visualization.

The ORFs for selected groups of phages (see the main text) were additionally annotated through HHblits searches against the Uniprot database clustered to 30% identity and the PDB database clustered to 70% identity (available at http://www.user.gwdg.de/~compbiol/data/hhsuite/databases/hhsuite_dbs/, accessed 02/2020) [40]. Genomes encoding a predicted reverse transcriptase (RT) were examined for the presence of repeats corresponding to a diversity-generating retroelement using DGRScan with default settings [41]. To identify repeats outside of the 10 kb RT-centered window (the default window of DGRScan), the template repeats were used as BLASTn queries against the encoding genome with the following parameters: -dust no -perc_identity 75 -qcov_hsp_perc 50 -ungapped - word_size 4.

### Fractional abundance of phage genomes in metagenomes

Dereplicated phage genomes from the NCBI Genbank database were combined with the dereplicated gut phages into a single database and indexed for read recruitment using Bowtie2 [42]. A collection of 1,256 human gut viromes were downloaded from the NCBI SRA using the SRA-toolkit (v. 2.10) and quality filtered with fastp (v. 0.20.1) [43]. The quality-filtered virome reads were recruited to the phage database using Bowtie2 with default parameters, except for the following additions “--no-unal --maxins 1000000”. The length-normalized fractional abundance of each phage genome in each virome was calculated as described previously [1].

### Host prediction from CRISPR-spacer matches

A database of CRISPR spacers was compiled from previous surveys of CRISPR-Cas systems [44, 45]. Each spacer was used as a BLASTN [46] query against the phage genomes, using word size of 8 and low complexity filtering disabled. A phage-host prediction was inferred if the spacer was 95% identical over 95% of its length to a phage sequence.

### Prediction of anti-CRISPR proteins

Identification of anti-CRISPR proteins (Acrs) was carried as out as previously described [47]. Briefly, each protein was assigned a score by the Acr prediction model ranging between 0 and 1, where a higher score corresponds to a higher likelihood of the protein being an Acr. The proteins were then clustered at 50% amino acid identity and considered a candidate Acr if they satisfied the following criteria: 1) received a mean score of 0.9 or above, 2) are present in a directon of 5 or fewer genes, 3) at least one of the directons encodes an HTH domain-containing protein and 4) the cluster does not produce a hit with an HHpred probability greater than 0.9 to any PDB or CDD database sequence [21].

## Results

### Identification of novel phage genomes from whole-community human gut metagenomes

The collection of 5,742 whole-community assembled metagenomes was searched for the presence of complete phage genomes. To limit the search space to putatively closed genomes, only ‘circular’ contigs with direct repeats (50 – 200 bp) at their termini (n = 95,663) were searched for open reading frames (ORFs) matching a known phage marker profile (i.e, the terminase large subunit, major capsid protein, or portal protein). In total, 3,738 contigs encode at least one ORF that passed the e-value and length cutoff criteria (Methods) (**Additional file 1**).

Dereplication at approximately 95% mean nucleotide identity reduced the number of phage marker-matching contigs to 1,886 (**Additional file 2**). A subset of 664 contigs encoded all three markers, 531 encoded two of the three and the remaining 691 possessed a single detectable marker (**Additional file 3**). The putative phage contigs had a median length of 44.9 kb which is consistent with the recent estimates of the median genome size of dsDNA phages [13]. To exclude any contaminating contigs (e.g., a plasmid harboring an integrated phage), each was assessed with ViralVerify [16] and Seeker [26], two bioinformatic tools trained to discriminate phage genomes from other sequences. These tools classified almost all the selected contigs as phages with varying levels of confidence except for several cases that, upon manual examination, were found to represent false negative classifications by these tools (**Additional file 2)**. Although we cannot rule out the possibility that some non-phage contigs were retained erroneously, the results collectively suggest that the set of circular marker-matching contigs predominantly consists of complete phage genomes.

To determine the host ranges of the phages, a database of CRISPR spacers from prokaryotic genomes was used to query the metagenomic phages for potential matches. In total, 553 (29%) of the dereplicated phage genomes were found to be targeted by at least one CRISPR-Cas system allowing host prediction (**Additional file 4**). The most common predicted hosts were *Firmicutes* (321 phages), followed by *Bacteroidetes* (143), *Actinobacteria* (43), *Proteobacteria* (41) and *Verrucomicrobia* (4).

Many phages have been found to encode Anti-CRISPR proteins (Acrs) to parry CRISPR-Cas defenses [48–51]. Given their function in counter-defense, Acrs evolve rapidly and show limited sequence similarity to experimentally characterized Acrs, making inference challenging [48]. However, a machine-learning based method has been recently developed that utilizes genomic context to identify candidate Acrs [47]. Application of this method showed that 41 phages, 16 of which were found to be targeted by a CRISPR-Cas system of their inferred host, encoded at least one candidate Acr, (**Additional file 5**). The highest-scoring Acrs belong to four phages that are targeted by *Bifidobacterium* CRISPR-Cas systems. All four phages are ≥ 97% identical over ≥ 90% of their length at the nucleotide level to uncharacterized prophages in cultured *Bifidobacterium* isolates **(Additional file 5)**, confirming their host-tropism assignment via CRISPR spacer-protospacer matches. In these phages, two candidate Acr-encoding genes lie between the large terminase subunit and integrase (**Additional file 6**). The localization of the Acr-encoding genes suggests they are expressed not only upon initial entry into the host cell and during lysogeny [52], but also upon transition to the lytic program to prevent cleavage of progeny phage genomes by CRISPR-Cas, as demonstrated experimentally in *Listeria*-infecting phages [53]. Transcription of Acrs is typically regulated by HTH domain-containing proteins termed Acr-associated proteins (Acas) [54]. Indeed, in the *Bifidobacterium* phages identified here, a short HTH domain-encoding ORF is located immediately downstream of the Acrs and can be predicted to regulate the expression of these two genes throughout the phage lifecycle. While these uncharacterized *Bifidobacterium* phages possess the characteristic features of Acr loci, the great majority of the phages identified in this work did not harbor any detectable Acrs yet were targeted by CRISPR-Cas (**Additional file 4)**. Some of these phages might encode distinct Acrs undetectable by the method we used that was trained on a collection of previously characterized Acrs, whereas others might employ alternative anti-CRISPR strategies.

### Taxonomic decomposition of the gut phages identifies previously unknown putative families

Phylogenetic trees were constructed for the large terminase subunit (TerL), major capsid protein (MCP) and portal protein encoded in each phage genome using an iterative approach to construct the underlying alignments [33]. The trees were constructed alongside reference proteins derived from phage genomes extracted from the NCBI GenBank database. Reflecting the set of protein profiles employed to identify the phage contigs, 1,480 (78%) genomes were assigned to the phylum *Uroviricota*, 360 (19%) to the *Phixviricota* and 46 (2%) to the *Loebvirae* (**Additional file 3**). The phylum *Phixviricota* includes *Escherichia coli* phage phiX174 that is used as a sequencing reagent; however, the 360 Phixviricota phages detected in this analysis do not include any sequences closely related to phiX174 (see Methods). The remaining analyses focus on the taxonomic decomposition of the phages that belong to *Uroviricota*, given that these contigs represent by far the largest fraction of recovered genomes.

The phylum *Uroviricota* is organized into a single class (*Caudoviricetes*) and order (*Caudovirales*), but a new order encompassing the crAss-like phages, a common and apparently most abundant group of phages in the human gut virome, is being proposed [55]. Our profiles recovered 141 phage genomes (dereplicated from 601 total genomes) that displayed phylogenetic relationship with the crAss-like phages and are the subject of a separate study (Yutin *et al.*, in press). The *Caudovirales* are presently organized into 9 families, but 3 of these (*Myoviridae, Podoviridae, Siphoviridae*) are expansive and demonstrably polyphyletic [56–58] and were thus not used for family-level taxonomic assignment although the remaining 6 families represent only a small fraction of the phages available in GenBank. The phylogenetic tree of TerL encoded by *Caudovirales* phages in our set reveals only 34 gut phages belong to one of these 6 ICTV-accepted families (Figure 1 and **Additional file 3**). The remainder of these unclassified phages are likely to found new families presently composed entirely of uncultured phages or belong to families with a cultured representative that have yet to be defined under the new multi-rank taxonomy of viruses.

**Figure 1:**
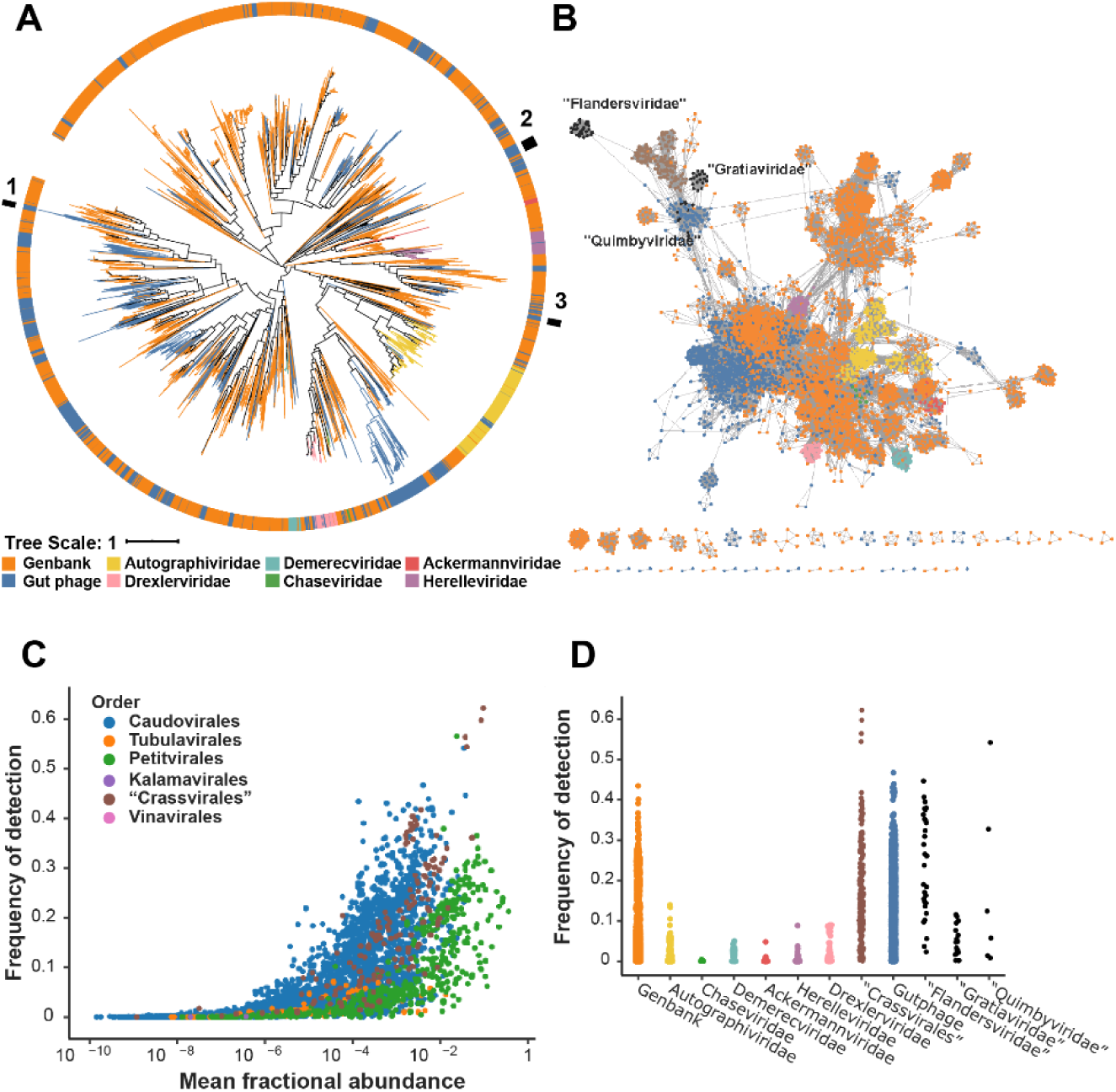
Three candidate families of Caudovirales phages discovered in human gut metagenomes. (A) Phylogenetic tree of the large terminase subunit encoded by Caudovirales phage genomes in GenBank (n = 3,931) and in gut metagenomes (n = 1,298). Branches are colored according to the current ICTV families, except for the Myoviridae, Podoviridae or Siphoviridae, which are in orange. The outermost ring indicates the location of candidate families proposed in this study: 1, “Quimbyviridae” phages; 2, “Flandersviridae” phages; 3, “Gratiaviridae” phages (see main text). (**B**) Gene sharing network of the Urovicota phages. Phage genomes identified in human gut metagenomes (blue nodes) were compared to phages in the GenBank database (colored as in Figure 1, with the addition of the crAss-like phages in brown and the new Caudovirales families proposed in this study in black). (**C**) Abundance of phages across human gut viromes. The x-axis depicts the fractional abundance of a given phage averaged across all viromes (n = 1,258); the y-axis is the fraction of viromes that a given phage recruits at least one read. Each phage genome (n = 7,888 total) is colored at the taxonomic level of Order (**C)** or Family (Uroviricota families only) (**D**). The raw data used to generate panel (A) are provided in **Additional file 14** and data for panels (C) and (D) are provided in **Additional file 15**.

### Selection of candidate families for comparative genomics

The taxonomic analysis based on phage hallmark proteins demonstrates that few phages in the human gut belong to a currently accepted ICTV-family. To prioritize candidate families for in-depth analysis, we next complemented the hallmark gene-based taxonomic analysis with whole-genome comparisons and abundance calculations of each phage relative to GenBank phages.

A gene-sharing network was constructed with the phages recovered from metagenomes and those deposited in GenBank. Edges are drawn between two viral genomes, represented as nodes, based on the number of ORFs that share significant sequence similarity [38]. Most of the metagenome-recovered phages bore multiple connections within the network to GenBank phages, in agreement with the manual curation of these contigs as genuine phage genomes (Figure 1B). However, two large groups of phages (tentatively labelled “Flandersviridae” and “Gratiaviridae”) were weakly connected to the larger network, reflecting disparate genome content. The divergence of the gene content of these phages from those of previously known phages and their distinct position in phylogenetic trees (Figure 1A and see below) indicate that they represent novel genera, and likely, new families.

To quantify the fractional abundance [1] of each phage in the human gut viral community, reads from a collection of 1,265 human gut viromes were competitively mapped against a database containing the metagenome-recovered and GenBank phages. The majority of the genomes do not recruit any reads (“detection”) from more than 2% of the viromes (Q1-Q3, 0-2% of viromes) (Figure 1C–D), consistent with the previously reported individuality of the human gut viromes [2, 7, 11]. A notable exception are the crAss-like phages [59] that recruit at least one read from about one-third of the viromes (Q1 - Q3, 9 - 28%), in agreement with previous reports of their cosmopolitan distribution [60, 61]. One uncharacterized *Caudovirales* genome was frequently observed in the collection of human gut viromes (54%, Figure 1C), suggesting that this phage is also cosmopolitan. To rule out the possibility that the observed frequency stemmed from non-specific read mapping to one or a few loci, rather than the complete genome, the coverage of sequencing reads across the genome (accession OMAC01000147.1) was examined. The broad coverage of this genome in the viromes confirms that its frequent detection is not an artifact, although several loci present in the reference sequence were absent in the viromes (**Additional file 7**). The exceptional detection of this uncharacterized phage (hereafter referred to as Quimbyvirus, after the character Mayor Quimby from the *Simpsons*) in the human gut viral community warrants its detailed examination.

Thus, three groups of phages were selected for in-depth analysis based on their distinct positions in the phylogenetic trees of the marker genes (all three groups), combined with divergent gene contents (“Flandersviridae” and “Gratiaviridae”) and high abundance in the human gut viral community (“Flandersviridae” and “Quimbyviridae”). A comparative genomic analysis of each candidate family is presented below, case-by-case.

### “Quimbyviridae” phages are abundant, hypervariable phages infecting Bacteroides

In the TerL phylogenetic tree, Quimbyvirus belongs to a group of phages whose closest characterized relatives include the *Vequintavirinae* and *Ounavirinae* subfamilies, under the now defunct *Myoviridae* family. To elucidate the taxonomic affiliation of Quimbyvirus, genomes from adjacent branches were examined (Figure 2). The median genome length of Quimby-like phages is 75.2 kb, close to the genome size of a branch basal to the Quimby-like branch, “group 4986” (72 kb), but smaller than the genomes of other phages in adjacent branches, *Ounavirinae* (88 kb) and *Vequintavirinae* (145 kb). Despite the similarity in genome size, phylogenetic reconstruction of the portal protein and MCP separate the Quimby-like phages from group 4986 (**Additional file 8**). Moreover, most Quimby-like phages encode a DnaG-family primase and DnaB-family helicase that are both absent in group 4986. However, in one branch of Quimby-like phages, the primase was lost from the replication module. The genomes of this branch encode a protein adjacent to the DnaB-family helicase with significant structural similarity to the winged helix-turn-helix domain of RepA (HHpred probability, 96.5) (Figure 2). RepA-family proteins mediate replication of plasmids by interacting with host DnaG primases [62], suggesting that the RepA-like protein coopts the host primase during replication, triggering the loss of the phage-encoded *dnaG* in this lineage. Consistent with a RepA-mediated episomal replication strategy, no integrase is identifiable in the genomes on this branch yet the phages encode numerous antirepressors, proteins involved in the lysis-lysogeny decision of temperate phages [63, 64]. The rest of the Quimby-like phages harbor a full-length integrase, indicating that these phages integrate into their host cell genome (**Figure**). Based on the topologies of the TerL, portal, MCP and DnaG trees, we propose that Quimby-like phages represent a novel taxonomic group at the family rank (henceforth, the “Quimbyviridae”). The potential differences in replication strategies (episomal vs. integrated) combined with the topologies of the phylogenetic trees of marker proteins suggest that “Quimbyviridae” splits into two distinct subfamilies.

**Figure 2:**
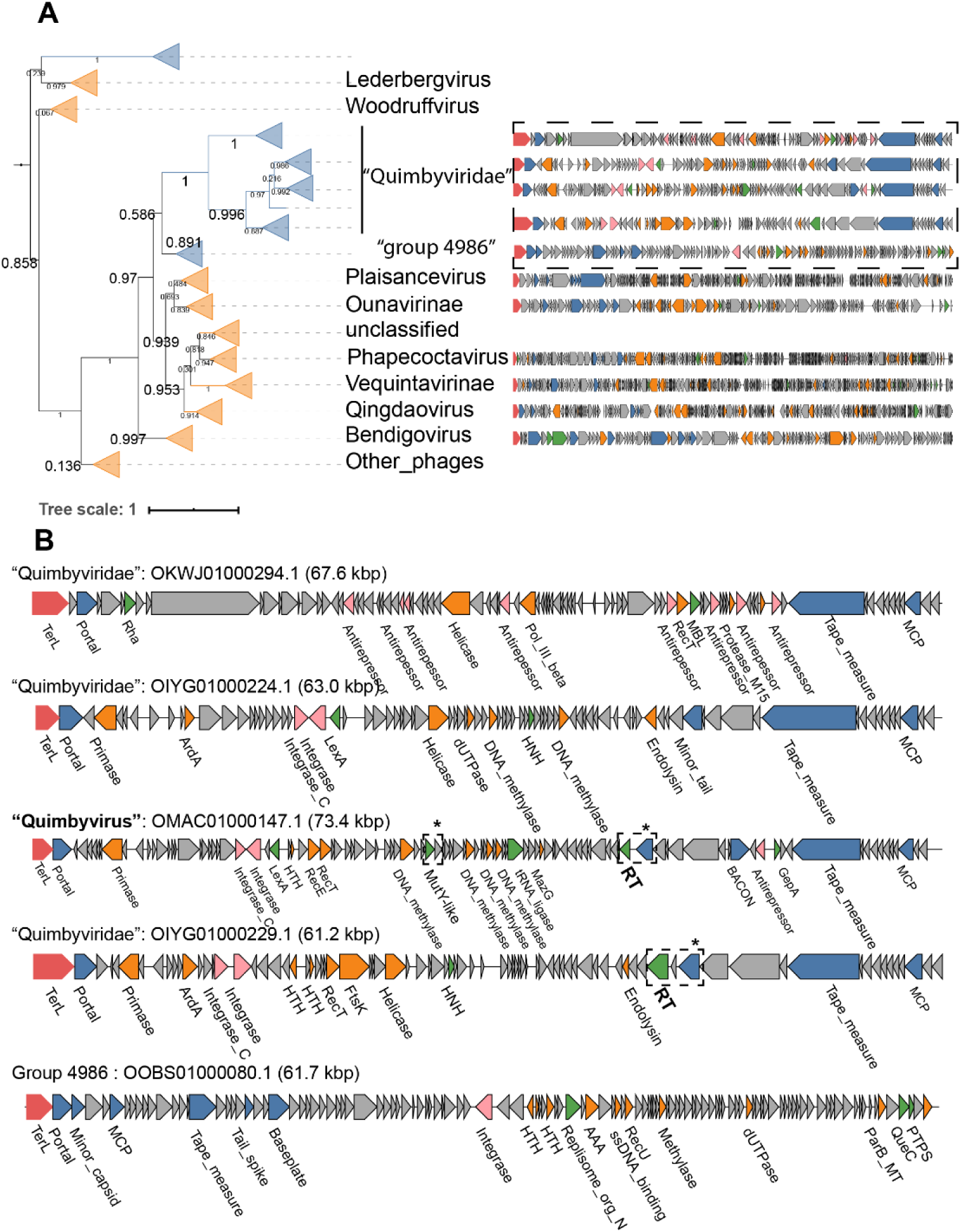
Phylogenetic tree of the large terminase subunit and genome maps of Quimby-like phages. (A) Individual genome maps of Quimby-like phages and ICTV classified phages are shown to the right of each branch. The ORFs are colored according to function: large terminase subunit (red), structural components (blue), DNA replication and repair (orange), lysogeny (pink), general function (green) and unknown (grey). **(B)** Expansion of four Quimby-like phages and a single gut phage genome from an adjacent branch (“group 4986”). The diversity-generating retroelement and hypervariable ORFs are highlighted with a dashed box and asterisk. The nucleotide scales differ between individual genome maps in both panels.

The Quimbyvirus genome aligns with a cryptic prophage of the bacterium *Bacteroides dorei* (CP011531.1), with 95% nucleotide sequence identity across 92% of its length, indicating that *B. dorei*, a common constituent of human gut microbiomes [65], carries a prophage closely related to Quimbyvirus. Inspection of the alignment shows that Quimbyvirus site-specifically integrates into the tRNA-Asp gene of *B. dorei*, a typical site of prophage integration [66]. The hosts of the other “Quimbyviridae” phages, determined through CRISPR-spacer analysis, include the *Prevotella*, *Bacteroides* and *Parabacteroides* genera within the phylum *Bacteroidetes* and the *Lachnospiraceae* within the phylum *Firmicutes*. In contrast, the hosts of group 4986 do not include any *Bacteroidetes*. The differences in the inferred host ranges support separating group 4986 from ‘Quimbyviridae” phages and suggests that group 4986 might represent a novel family, but these genomes were not investigated further.

Some of the “Quimbyviridae” phages harbor diversity-generating retroelements (DGRs), a cassette of genes that selectively mutate a short locus, known as the variable repeat, that is part of a C-type lectin or an immunoglobulin-like domain [67, 68]. Targeted mutation of these domains yields proteins with altered binding affinities and specificities [69]. The DGR cassette in *Bordetella* phage BPP-1 of the genus *Rauchvirus* is the only experimentally studied DGR system in a phage, where diversification of the C-type lectin domain-containing tail fiber gene enables adsorption to different host cell receptors [70]. In Quimbyvirus, the RT component of the DGR is encoded by overlapping ORFs in all three frames (ORFs 52-54), suggesting that the active RT is produced by two programmed frameshifts. Although overlapping ORFs and programmed frameshifts have been identified in in many compact tailed phage genomes [71–74], DGR RTs have thus far only been predicted to be encoded by a single ORF. To discern if the frameshifts render the RT inactive, the variable repeats were examined for adenine-specific substitutions, a hallmark of DGR-mediated variation [68]. The two variable repeats reside in ORF 47 and 80 of the Quimbyvirus genome which both encode proteins containing C-type lectin domains, the canonical target of DGRs [67] (Figure 2). Alignment of the variable repeats with their cognate template repeats from nearly identical Quimbyvirus genomes (> 95 % average nucleotide identity) allowed the detection of 22 adenine sites in the variable repeat exhibiting substitutions whereas all other bases were nearly perfectly conserved (**Additional file 9**). Collectively, these results suggest that the frameshifted RT possesses the selective infidelity that characterizes DGR-mediated hypervariation.

The first variable repeat resides in the C-terminus of ORF 51 that is located downstream of the tail fiber genes, suggesting that this gene codes for a structural component of the virion, similar to the hypervariable tail fiber of phage BPP-1 [70, 75]. The second DGR target locus is in ORF 84 that is distal to the phage structural gene module and is expressed from the opposite DNA strand, suggestive of a non-structural protein. The genomic neighborhood of ORF 80 includes genes coding for a nuclease, four methyltransferases and a tRNA ligase within 7 kb. The nuclease shows significant sequence and structural similarity to *E. coli mutY* (HHpred probability, 97.3, **Additional file 10**), a DNA glycosylase involved in base excision repair. The methyltransferases are most similar to adenine- and cytosine-modifying enzymes (HHpred probability 100 and 99.9, respectively, **Additional file 10**) that likely prevent cleavage by host restriction endonucleases. Similarly, the tRNA ligase might repair tRNAs cleaved by host anticodon endonucleases [76]. Overall, the adjacency of ORF 84 with defense- and counterdefense-related genes implies that this hypervariable phage protein plays a role in the phage-host conflicts; however, the exact functions of the DGR and hypervariable target proteins during the life cycle of “Quimbyviridae” phages remain to be investigated.

### “Flandersviridae” phages are common and abundant in whole-community metagenomes

Analysis of the phylogenetic trees of TerL identified a deep branch of 29 gut phages (dereplicated from 196 total genomes) that joins the family *Ackermannviridae* (Figure 3A). Annotation of the ORFs encoded by the 29 representative contigs demonstrated that the genomes are colinear, confirming that they belong to a cohesive group (Figure 3B). The cohesiveness of this group was confirmed by the gene-sharing network, where these genomes form a coherent cluster that has few connections to the larger network (Figure 1B), reflecting distant (if any) similarity between most of the proteins encoded by these phages and proteins of phages in GenBank. The median genome size of the phages in this group is 85.2 kb, compared to 157.7 kb among the *Ackermannviridae* phages. There is a conserved module of structural genes that encode the MCP, portal, sheath and baseplate proteins, TerL and the virion maturation proteinase. The presence of a contractile tail sheath indicates that these viruses possess contractile tails similar to those in the family *Ackermannviridae*, in agreement with the TerL phylogeny. Several of the genes within the structural block contain immunoglobulin-like or C-type lectin domains (e.g., BACON and GH5, respectively) which are predicted to play a role in adhesion of the virion to bacterial cells or host-associated mucosal glycans [77–80]. Downstream of the structural block is a module of genes involved in DNA replication that includes a DnaB-family helicase, DnaG-family primase and DNA polymerase I (PolA). The *polA* gene is widely distributed among dsDNA phages and therefore serves as a useful marker for delineating the diversity of phage replication modules [81]. Phylogenetic reconstruction of both *polA* and *dnaG* encoded by these phages confirmed their monophyly (**Additional file 11**). Following the replication module is an approximately 20 kb long locus containing ORFs that showed no detectable similarity to functionally characterized proteins. Two of the phages harbor matches to CRISPR spacers encoded by *Bacteroides* and *Parabacteroides* spp., indicating these bacteria serve as hosts. Based on the large terminase and *polA* phylogeny, colinearity of their genomes and differences from known phages in both genome size and content, we propose that these *Bacteroides-*infecting phages represent a novel taxonomic group, probably, with a family rank, hereafter, named “Flandersviridae” (after the region where some of the metagenomes were sampled).

**Figure 3:**
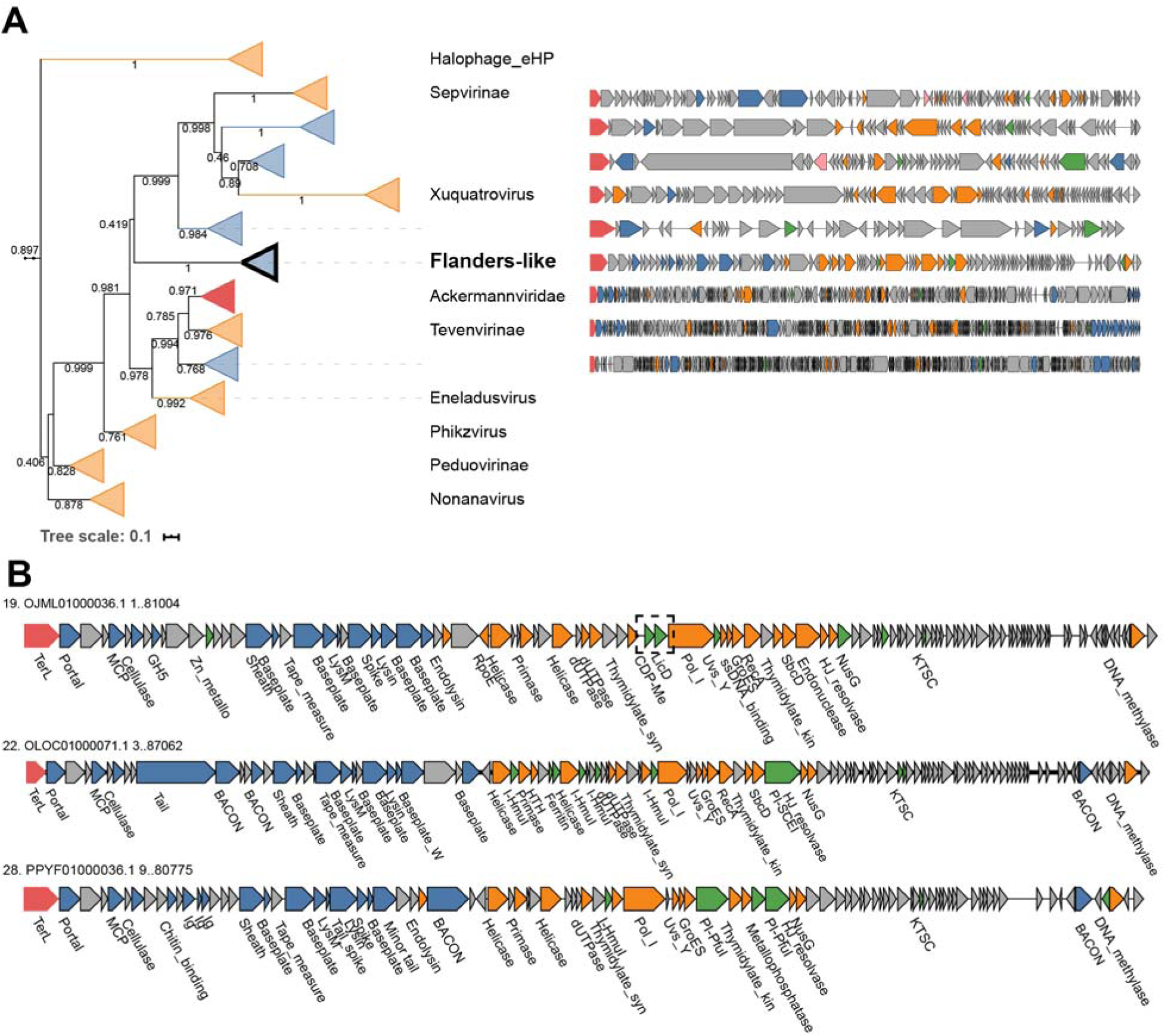
Phylogenetic tree of the large terminase subunit and complete genome maps for “Flandersviridae”. **(A)** Genome maps of members of the “Flandersviridae and selected ICTV-classified phages were constructed and colored as in Figure 3. **(B)** Genome maps of three genera from the “Flandersviridae” family. The dashed box highlights the insertion of licD- and ispD-family enzymes in the replication module of one “Flandersviridae” phage.

Although all members of the “Flandersviridae” are syntenic, some contain an insertion of two adjacent genes encoding nucleotidyltransferase superfamily enzymes within the DNA replication module. One enzyme belongs to the *ispD* family that is involved in the biosynthesis of isoprenoids [82, 83], and the other is a *licD* family enzyme that is responsible for the addition of phosphorylcholine to teichoic acids present in bacterial cell walls [84] (Figure 3). To our knowledge, neither of these enzymes has been reported in phages previously. Given that only some members of the “Flandersviridae” possess these genes, they are unlikely to perform essential functions in phage reproduction, and instead could be implicated in phage-host interactions. The *licD* family enzyme might modify teichoic acids to prevent superinfection by other phages, given that these polysaccharides serve as receptors for some phages to adsorb to the host cells [85]. The role of *ispD* is less clear because *ispD* family enzymes catalyze one step in the biosynthesis of isopentenyl pyrophosphate, a building block for a large variety of diverse isoprenoids [86]. Phages manipulate host metabolic networks including central carbon metabolism, nucleotide metabolism and translation [87]; the discovery of *ispD* present in the “Flandersviridae” phage genomes might add to this list the isoprenoid biosynthetic pathway.

Complete “Flandersviridae” phage genomes were recovered from 249 whole-community human gut metagenomic assemblies. Their frequent assembly into closed contigs suggests that these phages might persist in their host cells as extrachromosomal circular DNA molecules, similar to phage P1 [88]. However, neither genes involved in DNA partitioning nor lysis-lysogeny switches are readily identifiable in the “Flandersviridae” genomes. Thus, this group of phages might be obligately lytic although discerning the lifestyle of a phage from the genome sequence alone is challenging [89]. Regardless of their lifestyle, the frequent recovery of these phages from whole-community metagenomes implies that they are common members of the human gut virome. Indeed, the “Flandersviridae” phages reach similar detection frequency as the crAss-like phages (Figure 1D) although there are fewer Flanders-like phages in the database. Like the “Quimbyviridae”, the even coverage of sequencing reads across one “Flandersviridae” genome (accession OLOC0100071.1) confirms its detection is not artifactual (**Additional file 12**). The high fractional abundance and detection of Flanders-like phages in viromes generally agrees with their frequent assembly from whole community metagenomes although they were not the most abundant (see Discussion). Overall, Flanders-like phages represent a previously undetected phage group that is widely distributed in human gut viromes.

### “Gratiaviridae”, a putative novel family of phages infecting Bacteroides

A deeply branching cluster of 18 genomes (dereplicated from 45 total) is basal to the families Autographiviridae, Drexlerviridae and Chaseviridae on the TerL phylogenetic tree (Figure 4A). Although not commonly present in gut viromes (Figure 1D), the deep relationship between these contigs and established phage families prompts in depth genome analysis of these putative phages. All 18 genomes encode a DnaG-family primase and a DnaE-family polymerase, and phylogenetic reconstruction for these genes demonstrates monophyly of these phages, the sole exception being the *dnaE* gene of bacteriophage phiST, a marine *Cellulophaga*-infecting phage that belongs to the polyphyletic, currently defunct Siphoviridae family [90] (**Additional file 13**). The *dnaG* and *dnaE* genes are nested within a module of other replication-associated genes that include superfamily I and II helicases, SbcCD exonucleases and a RecA family ATPase (Figure 4B). The structural module is composed of genes that encode an MCP, capsid maturation protease, portal protein, baseplate proteins and a contractile tail sheath protein. Although these genomes are not strictly colinear as observed for the “Flandersviridae” phages, the overall similarity of the proteins encoded by these phages is apparent in the gene-sharing network where they form a coherent cluster that shares some edges with the crAss-like phages (Figure 1B). Similar to crAss-like phages, the predicted hosts suggested by CRISPR-spacer matches are the *Bacteroides* and *Parabacteroides* genera (**Additional file 4)**. Taken together, the phylogenetic and genomic organization of these phages indicate that they represent a new family, provisionally named “Gratiaviridae” (after the pioneering phage biologist Dr. Andre Gratia).

**Figure 4:**
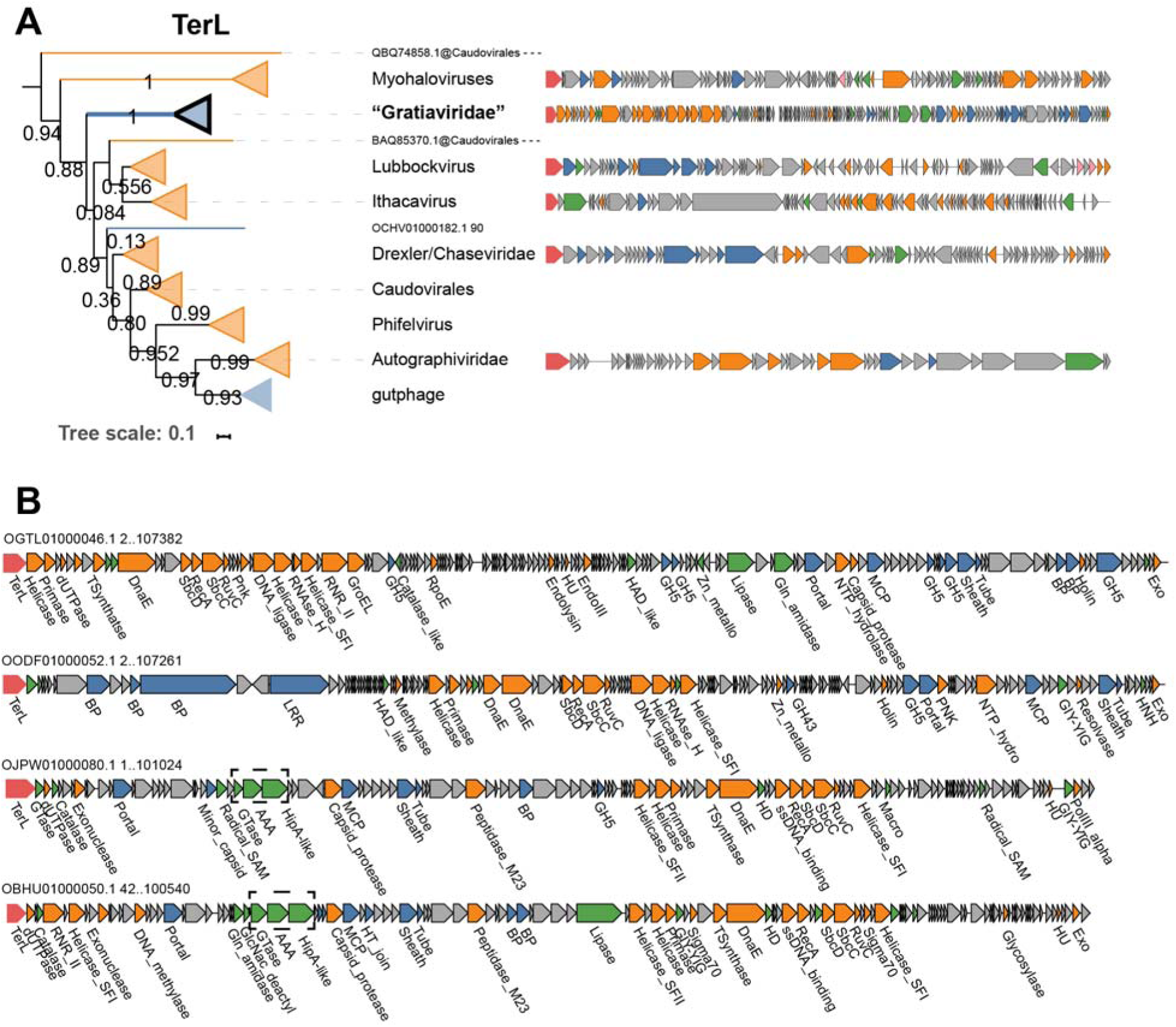
Phylogenetic tree of the large terminase subunit and genome maps of the “Gratiaviridae” phages. (A) Genome maps of ICTV-classified phages were constructed and colored as in Figure 3. (B) Genome maps of four genera from the Gratiaviridae family. The dashed box highlights a HipA-family kinase domain-containing protein, AAA-family ATPase and glycosyltransferase (see main text).

In addition to structural and replication proteins, “Gratiaviridae” phages encode several enzymes of the ferritin-like diiron-carboxylate superfamily. The ferritin-like enzymes encoded by these phages belong to two families, namely, DNA protecting proteins (DPS) and manganese-catalases. Manganese-catalases have not been documented in phage genomes and DPS-like enzymes have only been observed in seven *Lactobacillus-*infecting phages [91]. Both enzymes are involved in the tolerance of anaerobes to oxidative stress. Catalases detoxify hydrogen peroxide to oxygen and water, enhancing survival of anaerobic *Bacteroides* in the presence of oxygen [92]. DPS enzymes catalyze a reaction between oxygen and free iron to yield insoluble iron oxide, lowering the concentration of both intracellular oxygen and free iron levels that would otherwise react with hydrogen peroxide and produce a hydroxyl radical, the most toxic reactive oxygen species [93, 94]. “Gratiaviridae” phages might deploy catalase- and DPS-like enzymes during infection to enhance the tolerance of their strictly anaerobic *Bacteroides* hosts to oxidative damage. Notably, these enzymes were not restricted to the “Gratiaviridae” but could be identified in 196 (manganese catalase) and 36 (Dps) other phage genomes, including the “Flandersviridae”. The frequent identification of these enzymes in gut phage genomes underscores the importance of intracellular iron and reactive oxygen species concentration for productive infections in an anaerobic environment.

Five of the “Gratiaviridae” phages encode a protein containing a serine/threonine protein kinase domain with distant but significant sequence similarity to HipA family kinases (HHpred probability 99, **Additional file 10**). Whereas HipA family kinases are present in numerous, phylogenetically distinct bacterial genomes as the toxin component of a distinct variety of type II toxin-antitoxin systems [95, 96], there are only two characterized examples of protein kinases encoded by phages. The protein kinase of T7-like phages phosphorylates RNA polymerase and RNAse III early during infection as part of the takeover of the host cell transcriptional and translational machinery [97–99]. In contrast, the protein kinase of *E. coli* phage 933W is expressed during lysogeny and mediates abortive infection upon superinfection of the host cell by phage HK97 [100]. The HipA-like kinase is unlikely to function early during infection like the kinase of T7-like phages because, in all five “Gratiaviridae” phages, the kinase is encoded between the portal protein and MCP genes, which are expressed late during infection in numerous cultured phages [101, 102]. Instead, the kinase might confer immunity to heterotypic phage infection, analogous to the kinase encoded by 933W [100]. In support of an immunity-related role, an AAA-family ATPase and a glycosyltransferase are encoded immediately upstream of the kinase in all five phage genomes (Figure 4). Glycosyltransferases are encoded within capsular polysaccharide biosynthetic loci [103] and phase variation of the capsular polysaccharides confers immunity from phages that rely on these molecules for adsorption [104]. The specific roles of the HipA-family kinase, ATPase and glycosyltransferase are unknown but, collectively, these enzymes might modify host cell capsules, granting temporary immunity to heterotypic phage infection while the morphogenesis of “Gratiaviridae” progeny virions completes.

## Discussion

A search of human gut metagenomes identified 3,738 putative complete phage genomes. In an attempt to recover complete phage genomes, this analysis restricts the search space to metagenomic contigs with direct terminal repeats which are present at the termini of some phage genomes that consequently form circular assemblies [12]. Phages with different replication and DNA packaging strategies, such as members of the phylum *Preplasmiviricota* or *Escherichia* phage Mu [12], that lack direct repeats do not yield circular assemblies and thus were not detected here. As a result, the set of phage genomes recovered by this strategy is both biased and an underestimate. The results are also skewed towards smaller genomes that are more likely to assemble into a single contig although, in one metagenome, a 294 kb phage genome was identified (**Additional file 1)**. Phylogenetic and comparative genomic analyses suggest that this set of contigs includes many previously unnoticed lineages of phages, some, most likely, at the family rank.

Two groups of phages were selected for in-depth analysis based on their frequent recovery in metagenomes and viromes. Complete genomes of “Flandersviridae” and “Quimbyviridae” phages were assembled in 249 and 20 whole-community metagenomes, respectively. Yet, Quimbyvirus was more frequently detected in the viromes than any “Flandersviridae” phage (Figure 1D). The discrepancy can be attributed to several factors, including sampling bias, the greater number of “Flandersviridae” genomes in the reference database “diluting” the number of mapped reads per genome, or the presence of variable loci (e.g., the variable repeats of DGRs) that break contig assemblies [105]. Regardless, both groups encompass abundant members of the human gut virome. Predictably, the hosts of these phages include *Bacteroides spp.*, which are some of the most dominant bacterial taxa of the human gut [106] and serve as hosts for other common human gut phages [61, 107]. Much of the uncharacterized “dark matter” in these phage genomes is likely to be dedicated to preventing superinfection of the *Bacteroides* host cells by such phages and to counter the host defenses. Although in general defense systems in *Bacteroidetes* remain poorly characterized, most of the bacteria possess active CRISPR-Cas systems, and numerous CRISPR spacers targeting the phages analyzed here were detected. This implies that many if not most of the phages infecting *Bacteroidetes* would encode Acrs. However, the currently available prediction method that was trained on the sequences of previously identified Acrs detected putative Acrs only in a small minority of these phages. The remaining phages of *Bacteroidetes* might encode distinct Acrs or employ alternative anti-CRISPR strategies.

Several phage genera possess DGRs, including Quimbyvirus. Metagenomic surveys have shown that DGRs are enriched in the viruses that inhabit gastrointestinal environments [105, 108]. Combined with the isolation of DGR-carrying phages from human gut bacteria [109, 110], these observations reflect a prominent role of hypervariability underlying phage-host interactions in the gastrointestinal environment. Notably, Quimbyvirus and another DGR-carrying phage (Hankyphage, BK010646.1), lysogenize the same *Bacteroides* species and both phages are frequently detected in human gut viromes [110]. The commonalities aside, the Quimbyvirus DGR RT is encoded by three overlapping reading frames and targets two proteins, one in the structural module and one in a defense-related island. DGRs have been associated with putative defense and signaling systems in cyanobacterial and gammaproteobacterial genomes [111, 112], but beyond the presence of the C-type lectin fold, the hypervariable proteins possess few other recognizable domains that obfuscate their precise roles.

The third group analyzed in this study, the “Gratiaviridae”, is not abundant but occupies a deep position on the TerL tree relative to the *Autographiviridae*, *Chaseviridae* and *Drexlerviridae* families. Analysis of the “Gratiaviridae” genomes will facilitate the future organization of these families into higher taxonomic ranks, potentially, at the order level. Furthermore, analysis of the “Gratiaviridae” genomes demonstrated the presence of catalase- and DPS-family enzymes that arbitrate cellular responses to oxidative stress [113]. Oxygen concentrations vary along the length of the gastrointestinal tract, where the concentration is lower in the distal vs. proximal gut [114]. Oxygen also diffuses from tissues radially into the lumen [115] and, in combination with other factors, these gradients affect the structure and composition of the gastrointestinal microbiota [116]. The acquisition of oxygen detoxifying-enzymes by the “Gratiaviridae” and other gut phages signals a need to supplement their host cell’s tolerance to oxidative damage during infection, which might be especially important for cells that reside near the tissue surface where oxygen exposure is higher.

A unique feature of some “Gratiaviridae” phages is a HipA-family protein kinase. The T7-like phages (within the *Autographiviridae* family) and *Escherichia* phage 933W (currently unclassified at the family level) encode PKC-family protein kinases that function during host cell takeover and abortive infection, respectively [97, 100]. A third, CotH-family protein kinase domain is occasionally observed in phage genomes where it is fused to a hypervariable C-type lectin domain [67, 108], but these proteins are currently unstudied. The “Gratiaviridae” family phage recruited of a fourth family of protein kinases that, together with the phage encode glysosyltransferase, might modify the host cell envelope, contributing to the prevention of superinfection.

## Conclusions

In summary, comparative genomic analysis of the phages described here, along with the complementary analysis of crAss-like phages (Yutin *et. al*, in press), substantially increases the characterized diversity of phages, primarily, those infecting Bacteroidetes bacteria, which are major components of the human gut microbiome. These findings also expand the repertoire of phage gene functions, notably, by adding the isoprenoid metabolic pathway, catalase-like enzymes, HipA family protein kinases and hypervariable genes implicated in defense, and open multiple directions for experimental study.

## Declarations

### Ethics approval and consent to participate

Not applicable

### Consent for publication

Not applicable

### Availability of data and materials

All phage genomes are available in the NCBI GenBank database using the accession numbers listed in **Additional file 1**. The genomes, protein predictions and annotations are also provided at ftp://ftp.ncbi.nih.gov/pub/benlersm/gut_phages_2020/. Underlying alignments of marker protein sequences used to generate phylogenetic trees are provided in **Additional file 14.** Raw sequence data was downloaded from the NCBI SRA database using the accession numbers listed in **Additional file 15**.

### Competing interests

The authors declare that they have no competing interests.

### Funding

Funding for this project was provided by the Intramural Research Program of the National Institutes of Health (National Library of Medicine). The funding body had no role in the collection, analysis, and interpretation of data or writing of the manuscript.

### Authors contributions

NY, MR and DA analyzed the gut metagenomes; SS and AG executed the CRISPR analyses; SB analyzed the phage genomes and wrote the manuscript; PP and EVK conceived the project and wrote the manuscript. All authors read and approved the final manuscript.

## Acknowledgements

We are grateful for the assistance from Yuri Wolf with phylogenetic reconstruction of the phage hallmark genes and to Koonin group members for useful discussions. This work utilized the computational resources of the NIH HPC Biowulf cluster (http://hpc.nih.gov).

## Additional files

### Additional file 1

Clustering information for 3,378 gut phages identified in the study. Phage genomes were dereplicated at 95% identity over 80% of the contig length. The GenBank accession codes for each phage and the representative sequence is provided in columns 1 and 3, respectively (.csv).

### Additional file 2

Taxonomic information for 1,886 representative phage genomes. Each taxonomically classified phage genome was scored by ViralVerify, Seeker and vcontact2 and the assignments are provided in columns 9-10, 11 and 12-13, respectively (.csv).

### Additional file 3

Distribution of the marker profiles identified on each phage genome and pie chart representation of the gut phage taxonomy. (**A**) Histogram of marker proteins detected on each phage genome recovered from human gut metagenomes. Abbreviations are as follows: M, major capsid protein; P, portal, T, terminase large subunit. **(B)** Taxonomic assignments of the dereplicated contigs (n = 1,886), with the outermost ring corresponding to ICTV families (.pdf).

### Additional file 4

Host ranges inferred from CRISPR-spacer matches. The nucleotide coordinates of the protospacer are provided in columns 2 and 3. The sequence of the CRISPR spacer and protospacer are provided in columns 4 and 5. The taxonomic information of the host is listed in the subsequent columns (.csv).

### Additional file 5

Candidate anti-CRISPR (Acr) proteins encoded on each phage genome. Proteins predicted to be an Acr with high confidence (score > 0.9) and meeting all other heuristic criteria are listed along with their inferred host. For the *Bifidobacteria* phages (see the main text), the nucleotide coordinates of closely related prophages integrated in their host genome are provided in columns 6-8 (.csv)

### Additional file 6

Genome maps of predicted anti-CRISPR proteins (Acrs) in uncharacterized *Bifidobacteria* phages. Open reading frames are colored according to function: large terminase subunit (red), structural components (blue), replication (orange), integrase (pink), general function (green) and unknown (grey). The candidate Acrs are indicated with a dashed box (.pdf)

### Additional file 7

Coverage heatmap of a “Quimbyviridae” genome across human gut viromes. The coverage of the most abundant “Quimbyviridae” phage genome (accession OMAC01000147.1) is plotted as a heatmap, scaled from 0 – 100x per 100 bp window (.pdf).

### Additional file 8

Phylogenetic tree of the MCP, primase and portal proteins encoded by “Quimbyviridae” phages (.pdf).

### Additional file 9

Alignment of the template and variable repeats from the Quimbyvirus DGRs. The template repeat (TR) from Quimbyvirus is the first listed sequence and is followed by the variable repeats from either ORF80 (VR1) or ORF 47 (VR2) encoded in phage genomes that are nearly identical to Quimbyvirus (> 95 % average nucleotide identity). A total of 21 adenine residues (green) in the template repeat exhibit a least one substitution in a corresponding variable repeat (.pdf).

### Additional file 10

HHPred alignments of four Quimbyviridae proteins and one Gratiaviridae protein with their top-scoring templates, including a replication initiator protein, a cytosine-specific methyltransferase, an adenine-specific methyltransferase, a MutY nuclease and HipA kinase (.docx).

### Additional file 11

Phylogenetic tree of the polA and dnaG genes in Flanders-like phages. Branches composed of GenBank phages are colored in orange and branches of gut metagenomic phages in blue. Branches with sequences labelled as bacteria in the GenBank database, likely representing cryptic prophages, are colored in grey (.pdf).

### Additional file 12

Coverage heatmap of a “Flandersviridae” genome across human gut viromes. The coverage of the most abundant “Flandersviridae” phage genome (accession OLOC0100071.1) is plotted as a heatmap, scaled from 0 – 100x fold coverage per 100 bp window (.pdf).

### Additional file 13

Phylogenetic tree of the *dnaG* and *dnaE* genes in Gratiaviridae phages (.pdf).

### Additional file 14

Fasta-formatted alignments and Newick-formatted files underlying all trees generated in the study (.zip).

### Additional file 15

Abundance matrix of phage genomes in human gut viromes. The first column lists the GenBank accessions of each reference phage genome and the first row lists the SRA accessions for each virome. Values represent the length-normalized fractional abundance of each phage in each virome (see methods) (.zip).

## Notes

### Competing Interest Statement

The authors have declared no competing interest.

